# Secretoglobin family 1D member 2 (*SCGB1D2*) protein inhibits growth of *Borrelia burgdorferi* and affects susceptibility to Lyme disease

**DOI:** 10.1101/2022.05.27.493784

**Authors:** Satu Strausz, Grace Blacker, Sarah Galloway, Paige Hansen, Samuel E. Jones, Erin Sanders, Nasa Sinnott-Armstrong, FinnGen, Irving L. Weissman, Mark Daly, Tuomas Aivelo, Michal Caspi Tal, Hanna M. Ollila

**Author notes:** These authors contributed equally and either of them can list themselves second in their CV. **Author Contributions:** Designed, conducted, and analyzed genetic and expression data: S.S., S.E.J. N.S.A., M.D., T.A. and H.M.O. Designed, conducted, and analyzed experiments: G.B., S.G., and P.H. Designed and oversaw experiments: M.C.T Mentorship and intellectual contributions: E.S. and I.L.W. Wrote the manuscript: H.M.O, S.S., M.C.T, S.E.J., S.G., G.B, T.A.

## Abstract

Lyme disease is a tick-borne disease caused by bacteria of the genus *Borrelia*. The disease can initially manifest as an erythema migrans rash and, if able to evade the host immune defenses, can progress into a secondary stage chronic disease with debilitating physical or neurological manifestations^1,2^. The host factors that modulate susceptibility for Lyme disease have remained mostly unknown. Here we show a novel host defense mechanism against Lyme disease in humans. Using epidemiological and genetic data from FinnGen, we identify a common missense variant at the gene encoding for Secretoglobin family 1D member 2 (*SCGB1D2*) protein that increases the susceptibility for Lyme disease. The genetic variant changes proline at position 53 to leucine and is predicted as deleterious. Consequently, we validate the dysfunction of this protein variant using live *Borrelia burgdorferi (Bb)*. Recombinant reference *SCGB1D2* protein inhibits the growth of *Bb* twice as effectively as the recombinant *SCGB1D2* P53L deleterious missense variant. Together, these data suggest that *SCGB1D2* is a host defense factor present in the skin, sweat, and other secretions which protects against *Bb* infection. This finding provides a novel therapeutic avenue for drug development to prevent and treat Lyme disease.

## MAIN

Lyme disease (i.e., borreliosis) is an infectious disease caused by bacteria of the genus *Borrelia*, and transmitted by ticks. While most individuals have subsiding or short-term infection, others develop severe infection which requires intensive antibiotic treatment and may result in chronic illness^1,2^. Seasonality of the disease is well characterized in Finland, and an increasing number of patients with Lyme disease have emerged during recent decades^3,4^. Yet the biological risk factors and disease mechanisms for infection or severe illness are still only partially understood. Therefore, we performed a genome-wide association study (GWAS) on Lyme disease assessed by International Classification of Diseases (ICD)-9 and 10 codes and explored the phenotypic and genetic risk factors with the goal to finemap the most significant genetic associations and to understand underlying biology.

We utilized data from 342,499 individuals who have participated in the FinnGen project to estimate the effect of genetic variation for Lyme disease. Descriptive analysis showed that 44.3% of all participants were male in FinnGen. 5,248 (1.5%) of FinnGen participants had received a Lyme disease diagnosis between 1988 and 2021. 1,974 (37.6%) of the cases were male. The diagnoses were derived from ICD-codes in the Finnish national hospital and primary care registries (**Supplementary Table 1)**.

## RESULTS

### Human leukocyte antigen loci affect susceptibility to Lyme disease

We identified two genetic loci in a GWAS for Lyme disease (P < 5.0 × 10^−8^, **Figure 1 and Supplementary Table 2**). One of these was, perhaps unsurprisingly, located at the human leukocyte antigen (HLA) region on chromosome 6. The role of HLA is well-established to affect predisposition to infectious and autoimmune traits ^5,6^, and this association highlights the overall importance of the HLA in Lyme disease. Furthermore, the lead variant (rs9273375, P = 6.53×10^-11^) is located at the 3’ end of *HLA-DQB1* gene. In addition, finemapping of this association shows that the lead variant, rs9273375, likely reflects the signal from *HLA-DQB1*06:02* (dominant model’s P = 3.1×10^-10^, r^2^ = 0.888 between rs9273375 and *DQB1*06:02*, **Supple**^−^**mentary Table 3**). The specific alleles, and especially the *HLA-DQB1*06:02* allele or alleles in high linkage disequilibrium (LD) such as *HLA-DRB1*15:01*, have been previously associated with influenza-A infection ^7,8^, autoimmune diseases such as multiple sclerosis ^9^, and with type-1 narcolepsy^10^.

**Figure 1.**
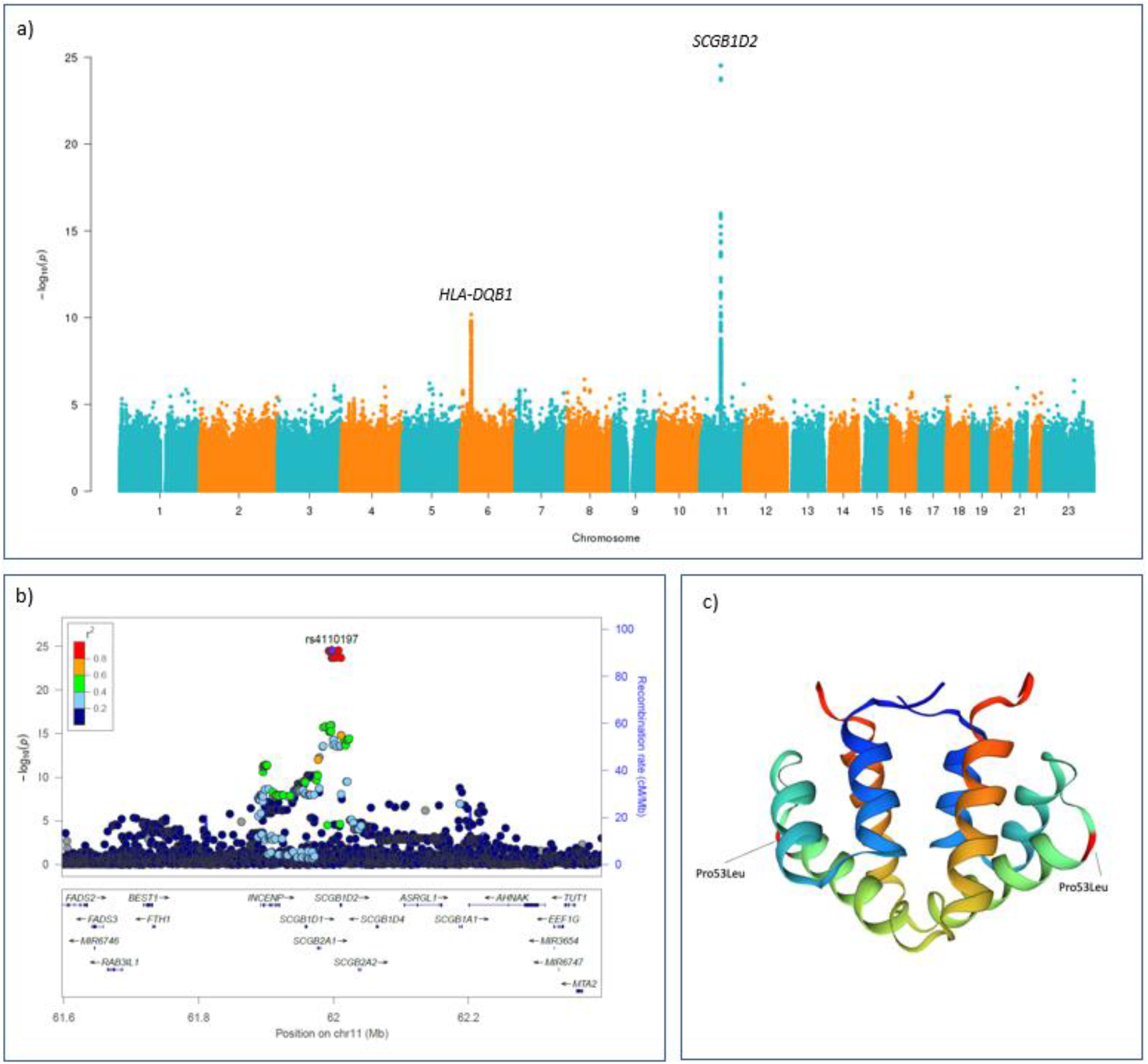
**a)** Manhattan plot for the genome-wide association study (GWAS) of Lyme disease (LD) including 5,248 LD cases and 337,251 controls. For each genetic variant, the x-axis shows chromosomal position, while y-axis shows the -log10(P) -value. The horizontal line indicates the genome-wide significance threshold of P=5.0x10^-8^. Two genetic loci were identified at a genome-wide significance level: SCGB1D2 and HLA-DQB1. **b)** Locus Zoom plot shows associated P-values on the −log_10_ scale on the vertical axis, and the chromosomal position along the horizontal axis. Purple diamond indicates single nucleotide polymorphism (SNP) at a locus with the strongest associated evidence. Linkage disequilibrium (LD, r^2^ values) between the lead SNP and the other SNPs are indicated by color. **c)** Schematic illustration for the protein structure of SCGB1D2 where a missense variant rs2232950 is causing an amino acid substitution from proline (Pro) to leucine (Leu). The structure is alpha-helical and forms an antiparallel dimer of the two monomers. There is a cavity which can accommodate small to medium sized ligands like steroids and phospholipids between the two dimers.

### Missense variants at *SCGB1D2* affect susceptibility to Lyme disease

While HLA association provides a proof of principle establishing Lyme disease as an infectious trait that is modified by genetic risk factors, the strongest and most compelling association was found at Secretoglobin family 1D member 2 locus (*SCGB1D2*, rs4110197, P = 2.95 × 10^-25^). rs4110197 is an intronic variant in *SCGB1D2* and finemapping^11^ of the locus indicated the causal variant is most likely among a set of eight single nucleotide polymorphisms (SNPs) including a common missense variant, rs2232950 (P = 1.82 × 10^-24^) with alternative allele frequency 40%, that is in high LD with the lead variant (r^2^ = 0.95). rs2232950 at position 158C>T of *SCGB1D2* results in a substitution from proline to leucine (Pro53Leu) and predicted deleterious by SIFT^12^ and Polyphen^13^ algorithms.

To understand the possible function of the variant, we examined its association in phenome-wide analysis (PheWAS) across 2,202 disease endpoints in FinnGen and publicly available GWAS data (**Supplementary Figure 1-2**). In addition to Lyme disease, we observed association with hospitalized spirochetal infections (Figure 2). As the spirochetal diseases include Lyme disease, we examined if the association was due to hospitalized Lyme disease patients and observed that the majority of hospitalized individuals had Lyme disease (N= 1,765 of 1,983 individuals with hospital level spirochetal infection).

**Figure 2.**
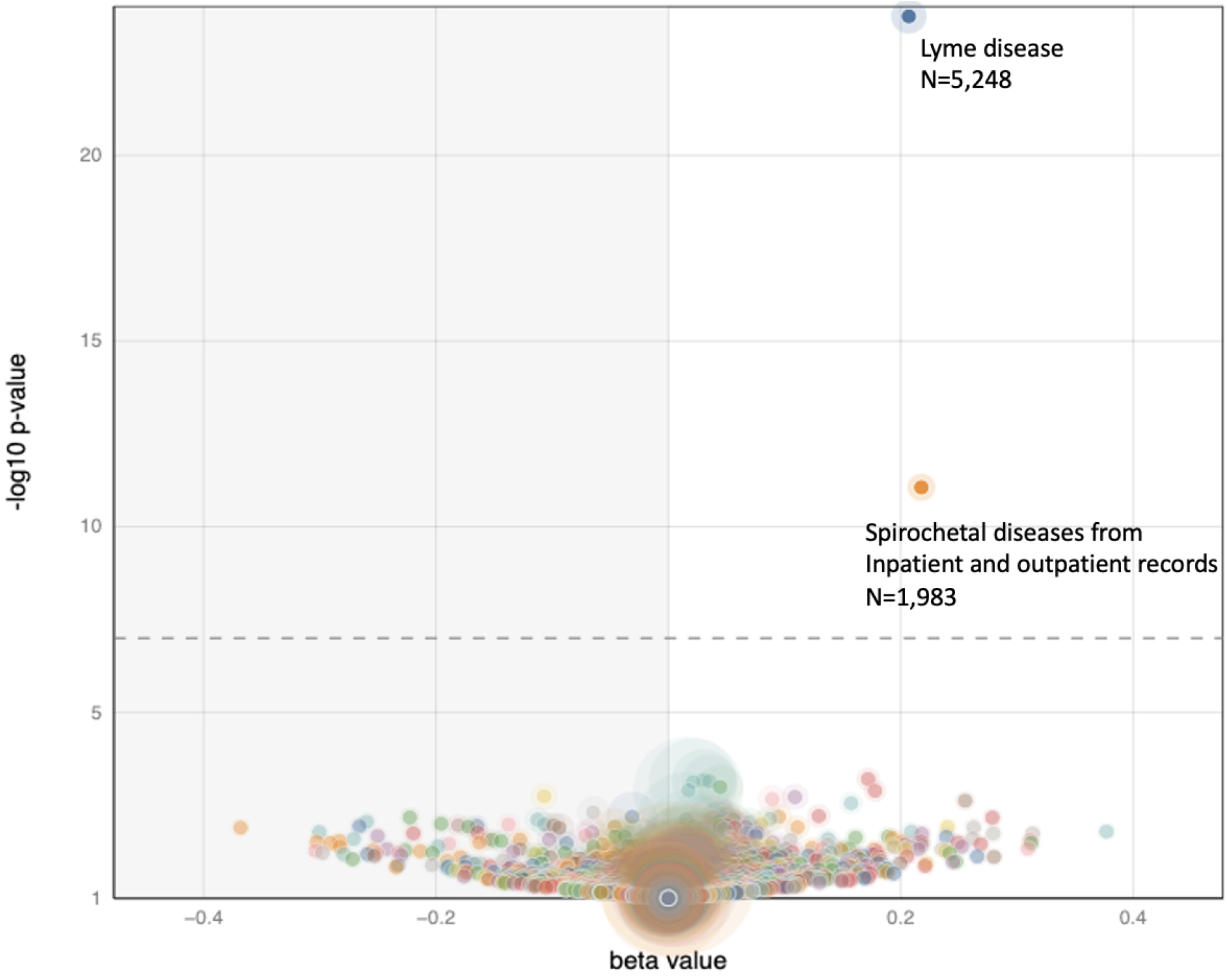
Volcano plot of Phenome-wide associations (PheWAS) from rs2232950 and 2,202 disease endpoints from FinnGen. Each point represents a trait. Vertical axis presents associated P-values at −log10 scale and the horizontal axis shows beta values. Other spirochetal diseases contain individuals who have been treated at hospital inpatient or outpatient clinics.

### *SCGB1D2* is expressed in the skin and secreted by sweat gland cells

*SCGB1D2* belongs to a Secretoglobin protein family in which all members, except for *SCGB1D2*, are found in upper airway tissues and are generally expressed by secretory tissues of barrier organs ^14,15^. We used data from The Genotype-Tissue Expression (GTEx) ^16^ release 8 to understand the tissue distribution of *SCGB1D2*. GTEx contains RNA expression samples from 948 donors across 54 tissues. The highest expression of *SCGB1D2* occurred in two skin tissues, sun exposed and not sun exposed skin (transcripts per million (TPM) = 258.2 and TPM = 163.2, respectively, **Supplementary Figure 3**). Expression was also observed in other secretory tissues including mammary tissue and the uterus. As skin is the first tissue for *Bb* exposure, we were interested in understanding the cell type distribution of *SCGB1D2* expression in the skin. Indeed, earlier studies have suggested that *SCGB1D2* expression may be even specific to sweat glands in the skin ^17,18^. Therefore, we visualized previously published single cell sequencing data across cell types that are observed in the skin: fibroblasts, keratinocytes, lymphatic endothelial cells, melanocytes, pericytes and smooth muscle cells, sweat gland cells, T cells and vascular endothelial cells ^17^. The visualization showed that *SCGB1D2* expression was specific to the sweat gland cells (**Supplementary Figure 4**) suggesting that *SCGB1D2* may be secreted on the skin as part of sweat. Antimicrobial peptides such as dermcidin, lysozyme, lactoferrin, psoriasin, cathelicidin and β-defensins have been previously discovered in human sweat ^19^, and raises the possibility that SCGB1D2 may have antimicrobial properties.

### *SCGB1D2* inhibits growth of *B. burgdorferi*

To investigate the impact of *SCGB1D2* protein encoded by the reference genotype on *Bb* growth we performed a *Bb* growth assay using recombinant reference *SCGB1D2*. We discovered that *SCGB1D2* inhibited the growth of *Bb* at 24h (ANOVA F (4, 18) = 5.77, P=0.0036) with significant growth inhibition at 8µg/mL and 16µg/mL concentrations (P=0.0091, P=0.0025 respectively, **Figure 3a)**. The growth inhibition was also significant at 72h (ANOVA F (4, 18) = 7.67, P=0.0009) time point at 8µg/mL and 16µg/mL concentrations (P=0.0137, P=0.0010, respectively, **Figure 3b**). Furthermore, despite an increase in the *Bb* count over time the growth inhibition was still notable at 72 hours suggesting that the effect is not transient (**Figure 3a-c)**. Furthermore, the growth inhibition was dose dependent at various *SCGB1D2* concentrations over time (**Figure 3c**).

**Figure 3.**
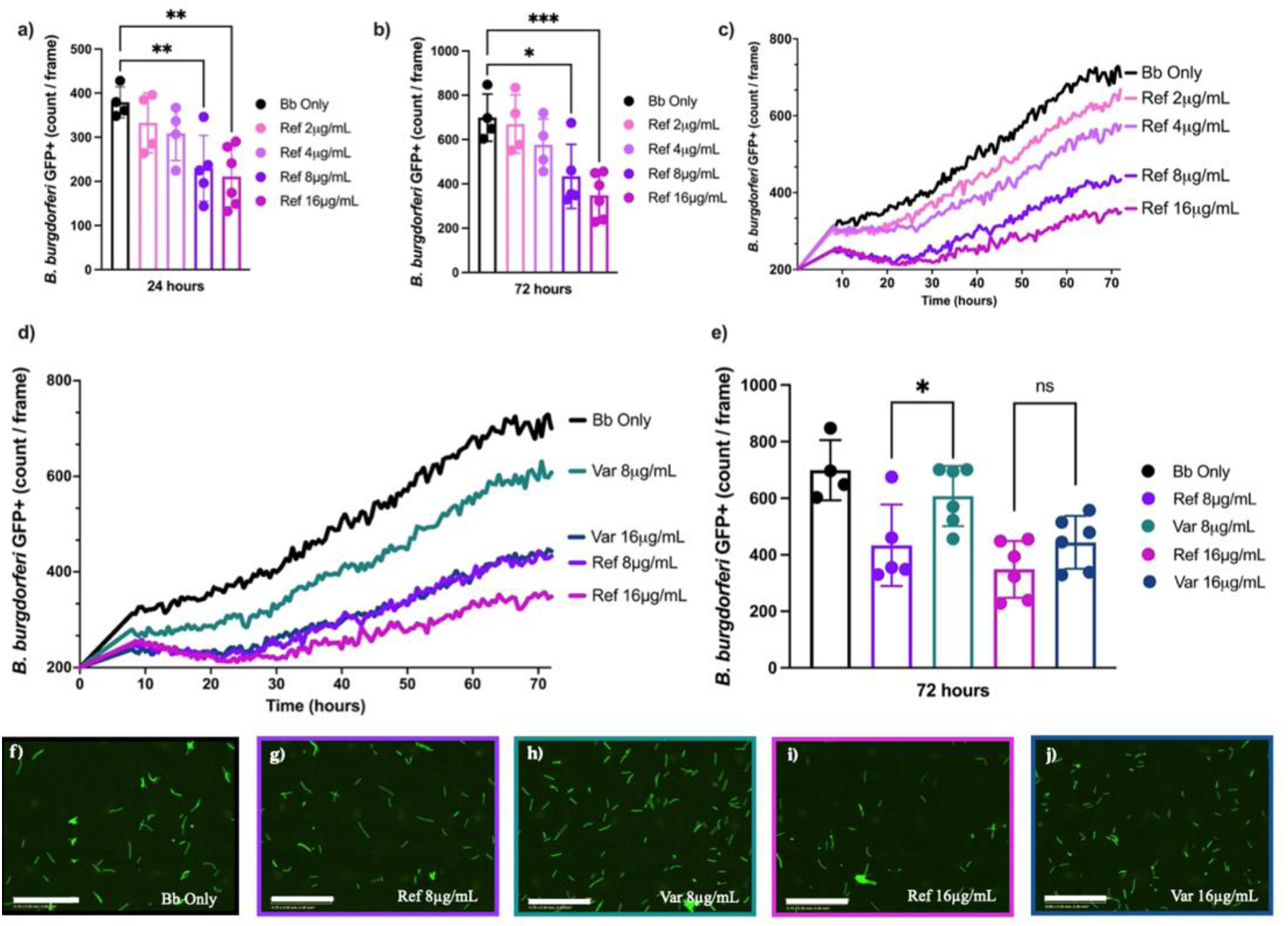
Borrelia burgdorferi (Bb) was incubated alone (black), or with either reference (Ref) or variant (Var) SCGB1D2 recombinant protein at 2µg/mL, 4µg/mL, 8µg/mL or 16µg/mL. IncuCyte analysis showing the count of green fluorescent protein (GFP)-expressing Bb per image with reference SCGB1D2 protein at 24h time point **a)** and at 72h time point **b)**. Count of GFP-expressing Bb spirochetes per image over time with all concentrations of reference SCGB1D2 protein **c)**. Count of GFP-expressing Bb spirochetes per image over time with SCGB1D2 reference and SCGB1D2 P53L recombinant proteins at 8µg/ml and 16 µg/mL concentration **d)**. Comparison of reference SCBG1D2 and variant SCGB1D2 P53L at 8µg/mL and 16µg/mL concentrations over time, and **e)** at 72 hours. Representative IncuCyte images for each treatment condition at 140 hours, scale bars at 200μm **(f-j)**. ***P=0.001, **P<0.01, * P<0.05 and ns, not significant.

We then tested whether the *SCGB1D2* P53L amino acid substitution encoded by the variant genotype might affect the *SCGB1D2* protein function or even have a different impact on *Bb* growth compared to the reference *SCGB1D2* protein without P53L substitution. To test this, we performed a *Bb* growth assay using recombinant reference *SCGB1D2* compared to recombinant variant *SCGB1D2* P53L. We discovered that approximately twice the amount of the variant *SCGB1D2* P53L was needed to achieve similar inhibition to *SCGB1D2* and that the variant *SCGB1D2* P53L inhibited Bb growth only at the highest concentration of 16µg/mL (**Figure 3d, Supplementary Figure 5**). Similarly, while the reference *SCGB1D2* overall inhibited *Bb* growth to a greater extent than *SCGB1D2* P53L variant at 8mg/mL, (P = 0.046, **Figure 3**) both reference *SCGB1D2* and *SCGB1D2* P53L variant were able to inhibit Bb growth at the highest concentration where the effect was possibly saturated (P = 0.119, **Figure 3 d-e**).

We observed that in the presence of 16 µg/mL of variant *SCGB1D2* P53L, and 8µg/mL or 16µg/mL of reference *SCGB1D2*, the *Bb* count reduces slightly from the starting amount to a trough around the 24-hour time point (**Figure 3d**). This may be indicative of bacterial killing by *SCGB1D2*, especially since this graphical trough is not observed in the *Bb* only condition or with 8µg/mL of variant *SCGB1D2* P53L.

In order to label dead cells and differentiate bacterial death from growth inhibition, propidium iodide (PI) was included in a repeat experiment of *Bb* growth inhibition. We observed a presence of dead spirochetes in the assay, and the cytotoxicity of prolonged PI exposure. Furthermore, a significant proportion of *Bb* were bound to PI compared to *Bb* conditions without PI, even in the absence of *SCGB1D2* proteins (**Supplementary Figure 6a**). To account for the increased PI binding, we examined the proportion of PI bound to *Bb* in the presence of either *SCGB1D2* reference or variant protein but did not observe significant difference of Bb death at 24 hours compared to Bb with PI control (**Supplementary Figure 6b**). Nevertheless, the graphical trends of *Bb* death in the presence of *SCGB1D2* protein were similar to observations of growth inhibition (**Figure 3)**. Specifically, 16µg/mL of reference *SCGB1D2*, which resulted in the most growth inhibition, also resulted in most PI binding (P = 0.0822). Further investigation is required to determine *SCGB1D2* mechanism of action against *Bb* and to differentiate bacterial killing from growth inhibition. Representative images of *Bb* from each treatment condition after 140 hours are also shown (**Figure 3f – 3j**).

## DISCUSSION

In this study, we report the first genetic variants that affect Lyme disease susceptibility. Most notably, we identify a novel association with a deleterious missense variant at the *SCGB1D2* gene. It is noteworthy that this *SCGB1D2* P53Lvariant appears quite specific for Lyme disease and has not been previously reported as associated with any other disease, phenotype, or infection. Using expression and single cell analysis, we observe that *SCBG1D2* has the highest expression in the skin and sweat gland cells. Furthermore, we characterized the function of *SCGB1D2* protein and show that recombinant *SCGB1D2* significantly inhibits *Bb* growth *in vitro*, and that around twice as much *SCGB1D2* variant protein is required to achieve the same level of *Bb* growth inhibition as reference *SCGB1D2*. These findings suggest that *SCGB1D2* is a restriction factor for *Bb* growth, and that *SCGB1D2* variant has a reduced antibacterial function against *Bb*. Overall, our results elucidate novel biology and indicate a mechanism by which a secretoglobin can provide protection against *Bb* infection in the skin. In the scope of infectious diseases, such disease-specific associations help teach us about specific host defenses that impact Lyme disease.

The current standard of care treatment for Lyme disease is bacteriostatic antibiotics such as amoxicillin or doxycycline, which restrict growth of bacteria similarly to *SCGB1D2* ^20^. Therefore, further investigating the prophylactic potential of *SCGB1D2* to prevent Lyme disease by restricting *Bb* growth and dissemination in early disease stages, may be of high interest, especially in individuals carrying the P53L mutation in *SCGB1D2*. Furthermore, it will be interesting to explore the potential of *SCGB1D2* applied topically as an intervention to reduce risk of Lyme disease. Similarly, *SCGB1D2* and similar molecules may open novel avenues for treating late-stage Lyme disease.

In the context of immune and infectious traits, it is nearly impossible to discuss risk for infection outside the scope of HLA. Also in our study, we identified a novel association with Lyme disease and *HLA-DQB1*06:02*. This finding highlights the overall impact that HLA alleles have on infectious and autoimmune traits and raises two interesting points. First, the balance between innate and adaptive immune responses in clearing *Bb* infection in the human host remains an active topic of research. HLA class II alleles such as *DQB1*06:02* are critical in modulating adaptive immune responses, and our findings suggest that adaptive immune responses, through primarily T-cell or B-cell mediated immunity, have an important role in Lyme disease. Second, both *HLA-DRB1*15:01* and *DQB1*06:02* have been previously implicated in brain autoimmune and infectious diseases such as multiple sclerosis, narcolepsy and Influenza-A^7,9,10^. *HLA-DRB1* has additionally been implicated in Lyme arthritis, and specifically the *HLA-DRB1*15:01* allele was found to be more common in antibiotic-refractory Lyme arthritis patients than antibiotic-responsive patients ^2^. Our findings provide yet another infectious disease trait that is associated with these same HLA alleles and raises the possibility that the same variants that contribute to infectious diseases also affect autoimmune and chronic disease traits in general.

Finally, our findings provide a compelling novel association with *SCGB1D2* protein as a restriction factor secreted in the skin and sweat. Additionally, they indicate a novel host defense mechanism against *Borrelia* infection and against Lyme disease. This finding provides a novel therapeutic avenue for drug development to prevent and treat Lyme disease.

## METHODS

### Data in FinnGen

FinnGen (www.finngen.fi/en) is a joint research project of the public and private sectors, launched in Finland in the autumn of 2017, that aims to genotype 500,000 Finns including prospective and retrospective epidemiological and disease-based cohorts as well as hospital biobank samples. FinnGen combines genome data with longitudinal health care registries using unique personal identification codes allowing data collection and follow-up even over the whole life span.

The FinnGen data release 8 is composed of 342,499 Finnish participants. The diagnosis of Lyme disease was based on ICD-codes (ICD-10: A69.2, ICD-9: 1048A), which were obtained from the Finnish national hospital and primary care registries including 5,248 individuals with Lyme disease and 337,251 controls. Main characteristics of the participants are outlined in Supplementary Table 1.

### Genotyping and imputation in FinnGen

FinnGen samples were genotyped with Illumina and Affymetrix chip arrays (Illumina Inc., San Diego, and Thermo Fisher Scientific, Santa Clara, CA, USA). Genotype calls were made with GenCall and zCall algorithms for Illumina and AxiomGT1 algorithm for Affymetrix data. Chip genotyping data produced with previous chip platforms and reference genome builds were lifted over to build version 38 (GRCh38/hg38) following the protocol described here: dx.doi.org/10.17504/protocols.io.nqtddwn.

In sample-wise quality control, individuals with ambiguous gender, high genotype missingness (>5%), excess heterozygosity (±4 standard deviation) and non-Finnish ancestry were excluded. In variant-wise quality control variants with high missingness (>2%), low Hardy-Weinberg equilibrium P-value (<1.0×10^-6^) and minor allele count < 3 were excluded. Prior imputation, chip genotyped samples were pre-phased with Eagle 2.3.5 ^21^with the default parameters, except the number of conditioning haplotypes was set to 20,000.

Genotype imputation was done with the population-specific SISu v4 reference panel. Variant call set was produced with Genomic analyses toolkit (GATK) HaplotypeCaller algorithm by following GATK best-practices for variant calling. Genotype-, sample- and variant-wise quality control was applied in an iterative manner by using the Hail framework v0.1 and the resulting high-quality whole genome sequenced data for 3,775 individuals were phased with Eagle 2.3.5. Post-imputation quality control involved excluding variants with INFO score < 0.7.

### Statistical methods

To compare the main characteristics of the participants and finemap HLA locus, a multivariate logistic regression model was used. A genome-wide association study was performed by Scalable and Accurate Implementation of Generalized mixed model (SAIGE) with a saddle point approximation to calibrate unbalanced case-control ratios ^22^. Analysis was adjusted for current age or the age at death, sex, genotyping chip, genetic relationship and first 10 principal components. To examine *SCGB1D2* locus and its genomic variation’s causality to Lyme disease in more detail, we finemapped this region utilizing the “Sum of Single Effects” -model, called *SuSiE* ^11^.

We performed a phenome-wide association analysis by retrieving association statistics for rs2232950 and all core phenotypes from FinnGen. We considered P-values significant if they passed Bonferroni correction for 2,202 tests corresponding to P-value 5.2×10^-5^.

We examined RNA expression across tissue types using GTEx v8 using the fully processed and normalized gene expression matrixes for each tissue ^16^. These same values are used for eQTL calculations by GTEx.

We used previously published single cell sequencing data ^17^ across cell types that are observed in the skin: fibroblasts, keratinocytes, lymphatic endothelial cells, melanocytes, pericytes and smooth muscle cells, sweat gland cells, T cells and vascular endothelial cells, and tested *SCGB1D2* expression in these cell types (Supplementary Table 4).

Data for functional analysis of *Bb* was collected by the IncuCyte® S3 and analyzed in GraphPad Prism v9.2.0. We conducted one-way ANOVA to explore the differences between groups followed by Dunnett’s post-hoc test to determine whether the tested groups were significantly different from the control group. We used independent T-tests in comparisons where there were only two groups.

### *B. burgdorferi* growth inhibition assay

B31A3-GFP (Green Fluorescent Protein) *Bb* was cultured at 37ºC in Barbour-Stonner-Kelly with 4-(2-hydroxyethyl)-1-piperazineethanesulfonic acid (HEPES) buffer media (BSK-H) complete with 6% rabbit serum (Millipore Sigma) and 1% Amphotericin B (Sigma-Aldrich). Bacterial concentration was determined by flow cytometry (Becton Dickinson LSRFortessa) such that 150,000 spirochetes per well of an ImageLock 96-well plate (Sartorius) were incubated with either 8µg/mL or 16µg/mL of variant (TP607036;MKLSVCLLLVTLALCCYQANAEFCPALVSELLDFFFISEPLFKLSLAKFDAPLEAVAAKLGVKRCTDQMSLQKRSLIAEVLVKILKKCSV) or reference (TP607035; MKLSVCLLLVTLALCCYQANAEFCPALVSELLDFFFISEPLFKLSLAKFDAPPEAVAAKLGVKRCTDQMSLQKRSLIAEVLVKILKKCSV) *SCGB1D2* recombinant protein (Origene Technologies) in a total well volume of 150uL. Incubation at 37ºC and 5% CO2 occurred inside of an IncuCyte® S3 (Sartorius) to measure real-time fluorescent intensity and capture phase images over time. Images were acquired using a 20x objective at 300-ms exposure per field of view. In order to limit false positive background fluorescence, threshold values to determine GFP^+^ events were set such that only *Bb* in phase and overlapped with green fluorescence events were counted.

A repeat *Bb* growth inhibition assay was performed as previously described, with the addition of 1.5μL propidium iodide (Millipore Sigma) per well prior to incubation. After 24 hours of incubation, samples were fixed in 4% paraformaldehyde, resuspended in flow cytometry buffer (2% FBS, 1mmol EDTA, in PBS), and analyzed by flow cytometry.

### Ethics statement

Patients and control subjects in FinnGen provided informed consent for biobank research, based on the Finnish Biobank Act. Alternatively, separate research cohorts, collected prior the Finnish Biobank Act came into effect (in September 2013) and start of FinnGen (August 2017), were collected based on study-specific consents and later transferred to the Finnish biobanks after approval by Fimea (Finnish Medicines Agency), the National Supervisory Authority for Welfare and Health. Recruitment protocols followed the biobank protocols approved by Fimea. The Coordinating Ethics Committee of the Hospital District of Helsinki and Uusimaa (HUS) statement number for the FinnGen study is Nr HUS/990/2017.

The FinnGen study is approved by Finnish Institute for Health and Welfare (permit numbers: THL/2031/6.02.00/2017, THL/1101/5.05.00/2017, THL/341/6.02.00/2018, THL/2222/6.02.00/2018, THL/283/6.02.00/2019, THL/1721/5.05.00/2019 and THL/1524/5.05.00/2020), Digital and population data service agency (permit numbers: VRK43431/2017-3, VRK/6909/2018-3, VRK/4415/2019-3), the Social Insurance Institution (permit numbers: KELA 58/522/2017, KELA 131/522/2018, KELA 70/522/2019, KELA 98/522/2019, KELA 134/522/2019, KELA 138/522/2019, KELA 2/522/2020, KELA 16/522/2020), Findata permit numbers THL/2364/14.02/2020, THL/4055/14.06.00/2020,,THL/3433/14.06.00/2020, THL/4432/14.06/2020, THL/5189/14.06/2020, THL/5894/14.06.00/2020, THL/6619/14.06.00/2020, THL/209/14.06.00/2021, THL/688/14.06.00/2021, THL/1284/14.06.00/2021, THL/1965/14.06.00/2021, THL/5546/14.02.00/2020, THL/2658/14.06.00/2021, THL/4235/14.06.00/2021 and Statistics Finland (permit numbers: TK-53-1041-17 and TK/143/07.03.00/2020 (earlier TK-53-90-20) TK/1735/07.03.00/2021).

The Biobank Access Decisions for FinnGen samples and data utilized in FinnGen Data Freeze 8 include: THL Biobank BB2017_55, BB2017_111, BB2018_19, BB_2018_34, BB_2018_67, BB2018_71, BB2019_7, BB2019_8, BB2019_26, BB2020_1, Finnish Red Cross Blood Service Biobank 7.12.2017, Helsinki Biobank HUS/359/2017, Auria Biobank AB17-5154 and amendment #1 (August 17 2020), AB20-5926 and amendment #1 (April 23 2020), Biobank Borealis of Northern Finland_2017_1013, Biobank of Eastern Finland 1186/2018 and amendment 22 § /2020, Finnish Clinical Biobank Tampere MH0004 and amendments (21.02.2020 & 06.10.2020), Central Finland Biobank 1-2017, and Terveystalo Biobank STB 2018001.

## Supporting information

FinnGen author banner

## Acknowledgements

We want to acknowledge the participants and investigators of FinnGen study. The FinnGen project is funded by two grants from Business Finland (HUS 4685/31/2016 and UH 4386/31/2016) and the following industry partners: AbbVie Inc., AstraZeneca UK Ltd, Biogen MA Inc., Bristol Myers Squibb (and Celgene Corporation & Celgene International II Sàrl), Genentech Inc., Merck Sharp & Dohme LCC, Pfizer Inc., GlaxoSmithKline Intellectual Property Development Ltd., Sanofi US Services Inc., Maze Therapeutics Inc., Janssen Biotech Inc, Novartis AG, and Boehringer Ingelheim International GmbH. Following biobanks are acknowledged for delivering biobank samples to FinnGen: Auria Biobank (www.auria.fi/biopankki), THL Biobank (www.thl.fi/biobank), Helsinki Biobank (www.helsinginbiopankki.fi), Biobank Borealis of Northern Finland (https://www.ppshp.fi/Tutkimus-ja-opetus/Biopankki/Pages/Biobank-Borealis-briefly-in-English.aspx), Finnish Clinical Biobank Tampere (www.tays.fi/en-US/Research_and_development/Finnish_Clinical_Biobank_Tampere), Biobank of Eastern Finland (www.ita-suomenbiopankki.fi/en), Central Finland Biobank (www.ksshp.fi/fi-FI/Potilaalle/Biopankki), Finnish Red Cross Blood Service Biobank (www.veripalvelu.fi/verenluovutus/biopankkitoiminta) and Terveystalo Biobank (www.terveystalo.com/fi/Yritystietoa/Terveystalo-Biopankki/Biopankki/). All Finnish Biobanks are members of BBMRI.fi infrastructure (www.bbmri.fi). Finnish Biobank Cooperative -FINBB (https://finbb.fi/) is the coordinator of BBMRI-ERIC operations in Finland. The Finnish biobank data can be accessed through the Fingenious® services (https://site.fingenious.fi/en/) managed by FINBB.

## References

1 Steere, A. C., Schoen, R. T. & Taylor, E. The clinical evolution of Lyme arthritis. Ann Intern Med 107, 725–731, doi:10.7326/0003-4819-107-5-725 (1987).

2 Wormser, G. P. et al. The clinical assessment, treatment, and prevention of lyme disease, human granulocytic anaplasmosis, and babesiosis: clinical practice guidelines by the Infectious Diseases Society of America. Clin Infect Dis 43, 1089–1134, doi:10.1086/508667 (2006).

3 Sajanti, E. et al. Lyme Borreliosis in Finland, 1995-2014. Emerg Infect Dis 23, 1282–1288, doi:10.3201/eid2308.161273 (2017).

4 Rizzoli, A. et al. Lyme borreliosis in Europe. Euro Surveill 16 (2011).

5 Tian, C. et al. Genome-wide association and HLA region fine-mapping studies identify susceptibility loci for multiple common infections. Nat Commun 8, 599, doi:10.1038/s41467-017-00257-5 (2017).

6 Lenz, T. L. et al. Widespread non-additive and interaction effects within HLA loci modulate the risk of autoimmune diseases. Nat Genet 47, 1085–1090, doi:10.1038/ng.3379 (2015).

7 Hammer, C. et al. Amino Acid Variation in HLA Class II Proteins Is a Major Determinant of Humoral Response to Common Viruses. Am J Hum Genet 97, 738–743, doi:10.1016/j.ajhg.2015.09.008 (2015).

8 Narwaney, K. J. et al. Association of HLA class II genes with clinical hyporesponsiveness to trivalent inactivated influenza vaccine in children. Vaccine 31, 1123–1128, doi:10.1016/j.vaccine.2012.12.026 (2013).

9 International Multiple Sclerosis Genetics, C. et al. Genetic risk and a primary role for cell-mediated immune mechanisms in multiple sclerosis. Nature 476, 214–219, doi:10.1038/nature10251 (2011).

10 Mignot, E. et al. Complex HLA-DR and -DQ interactions confer risk of narcolepsy-cataplexy in three ethnic groups. Am J Hum Genet 68, 686–699, doi:10.1086/318799 (2001).

11 Wang G, S. A., Carbonetto P, Stephens M. A Simple New Approach to Variable Selection in Regression, with Application to Genetic Fine Mapping. Journal of the Royal Statistical Society: Series B (Statistical Methodology). 82, 1273–1300 (2020).

12 Ng, P. C. & Henikoff, S. SIFT: Predicting amino acid changes that affect protein function. Nucleic Acids Res 31, 3812–3814, doi:10.1093/nar/gkg509 (2003).

13 Adzhubei, I. A. et al. A method and server for predicting damaging missense mutations. Nat Methods 7, 248–249, doi:10.1038/nmeth0410-248 (2010).

14 Jackson, B. C. et al. Update of the human secretoglobin (SCGB) gene superfamily and an example of ‘evolutionary bloom’ of androgen-binding protein genes within the mouse Scgb gene superfamily. Hum Genomics 5, 691–702, doi:10.1186/1479-7364-5-6-691 (2011).

15 Lu, X. et al. The cytokine-driven regulation of secretoglobins in normal human upper airway and their expression, particularly that of uteroglobin-related protein 1, in chronic rhinosinusitis. Respir Res 12, 28, doi:10.1186/1465-9921-12-28 (2011).

16 Consortium, G. T. The Genotype-Tissue Expression (GTEx) project. Nat Genet 45, 580–585, doi:10.1038/ng.2653 (2013).

17 He, H. et al. Single-cell transcriptome analysis of human skin identifies novel fibroblast subpopulation and enrichment of immune subsets in atopic dermatitis. J Allergy Clin Immunol 145, 1615–1628, doi:10.1016/j.jaci.2020.01.042 (2020).

18 Na, C. H. et al. Integrated Transcriptomic and Proteomic Analysis of Human Eccrine Sweat Glands Identifies Missing and Novel Proteins. Mol Cell Proteomics 18, 1382–1395, doi:10.1074/mcp.RA118.001101 (2019).

19 Csosz, E., Emri, G., Kallo, G., Tsaprailis, G. & Tozser, J. Highly abundant defense proteins in human sweat as revealed by targeted proteomics and label-free quantification mass spectrometry. J Eur Acad Dermatol Venereol 29, 2024–2031, doi:10.1111/jdv.13221 (2015).

20 Lantos, P. M. et al. Clinical Practice Guidelines by the Infectious Diseases Society of America (IDSA), American Academy of Neurology (AAN), and American College of Rheumatology (ACR): 2020 Guidelines for the Prevention, Diagnosis and Treatment of Lyme Disease. Clin Infect Dis 72, 1–8, doi:10.1093/cid/ciab049 (2021).

21 Loh, P. R. et al. Reference-based phasing using the Haplotype Reference Consortium panel. Nat Genet 48, 1443–1448, doi:10.1038/ng.3679 (2016).

22 Zhou, W. et al. Efficiently controlling for case-control imbalance and sample relatedness in large-scale genetic association studies. Nat Genet 50, 1335–1341, doi:10.1038/s41588-018-0184-y (2018).

