## Supplementary material for "Secretoglobin family 1D member 2 (*SCGB1D2*) protein inhibits growth of *Borrelia burgdorferi* and affects susceptibility to Lyme disease"

The purpose of this document is to provide information on the cohorts, choices for analytical methods and findings that are not included in the main paper, or where additional details may be useful.

**Contents**

1. Demographic characteristics
2. Analysis of individual genetic variants
3. Understanding of genetic association
    a. *HLA* locus
    b. *SCGB1D2* locus
4. PheWAS analysis
    a. Comparison with publicly available traits
    b. GWAS for syphilis
5. Expression profiling
    a. GTEx
    b. Single cell analysis
6. Functional analyses of *Borrelia burgdorferi*

### 1. Demographic characteristics

For our study, we used the FinnGen cohort, which combines genomic data with national health registries.

We defined Lyme disease by extracting International Classification of Diseases (ICD)-9 (1048A) and ICD-10 (A69.2) codes from hospital inpatient, hospital outpatient and primary outpatient health registries. Our data consisted of 5,248 individuals with Lyme disease and 337,251 controls. Of all patients with Lyme disease diagnosis 37.6% were male and the mean age was 65.1 years. The mean age at the first Lyme disease diagnosis was 60.5 years (Supplementary Table 1).

**Supplementary Table 1.** *Demographic and diagnosis information in individuals diagnosed with Lyme disease in FinnGen*

| | Lyme | Non-Lyme | OR [95% CI] | P Lyme vs. Non-lyme |
|---|---|---|---|---|
| **N ICD-10 A69.2** | 5,189 | | | |
| **N ICD-9 1048A** | 59 | | | |
| **N total** | 5,248 (1.5%) | 337,251 | | |
| **Sex (male)** | 1,974 (37.6%) | 149,646 (44.4%) | 1.50 [1.41-1.58] | $< 2\times10^{-16}$ |
| **Age (mean, SD)** | 65.1 (14.3) | 59.0 (18.0) | 1.02 [1.021-1.024] | $< 2\times10^{-16}$ |
| **Age at first diagnosis for Lyme (mean, SD)** | 60.5 (15.1) | | | |

*Comparison between individuals with and without Lyme disease diagnosis in FinnGen. ICD=International Classification of Diseases, OR=odds ratio, CI=confidence interval, SD=standard deviation.*

### 2. Analysis of individual genetic variants

To study genetics behind Lyme disease we analyzed a total of 342,499 samples from the FinnGen Data Freeze 8 with 5,248 individuals with Lyme disease diagnosis. For the GWAS we utilized Scalable and accurate implementation of generalized mixed model (SAIGE) [22], which uses saddle point approximation to calibrate unbalanced case-control ratio. In addition, this method reduces the risk for type 1 error. Our GWAS analysis was adjusted for current age or the age at death, sex, genotyping chip, genetic relationship and first the 10 principal components.

The GWAS revealed two genome-wide significant signals ($P < 5.0\times10^{-8}$). The characteristics of these loci are presented in **Supplementary Table 2**. Our analysis pointed to a genome-wide signal in the HLA-region (rs9273375) located at the 3' end of *HLA-DQB1*. In addition, we observed the strongest association in *SCGB1D2* where the lead intronic variant (rs4110197) was found to be in high LD with a missense variant (rs2232950). Curiously, this variant causes amino acid change from proline to leucine, and this change is predicted deleterious by several databases.

**Supplementary Table 2.** *Genome-wide lead variants for Lyme disease*

| CHR | rsid | REF | ALT | AF cases | AF controls | Finnish enrichment | OR [95% CI] |
|---|---|---|---|---|---|---|---|
| **6** | rs9273375 | G | C | 0.83 | 0.85 | 1.01 | 0.83 [0.79-0.88] |
| **11** | rs4110197 | A | G | 0.45 | 0.40 | 1.27 | 1.24 [1.19-1.29] |

*Characterization of two genome-wide significant Lyme disease loci. Effect sizes and allele frequencies are reported in terms of alternative allele (ALT). Finnish enrichment is computed using the Genome Aggregation Database (gnomAD) data comparing Finnish individuals to other European populations. CHR=chromosome, REF=reference allele, AF=allele frequency, OR=odds ratio, CI=confidence interval.*

### 3. Understanding of genetic associations

#### a) HLA locus

HLA has a strong and established role in human immune defense. However, its contribution to Lyme disease has not been previously thoroughly explored. Our GWAS results showed a genome-wide significant HLA-locus, and to study this finding in more detail we finemapped this region. We computed association statistics with each HLA-allele from the *HLA-A*, *HLA-B*, *HLA-C*, *HLA-DRB1*, *HLA-DQA1*, *HLA-DQB1*, *HLA-DPA1* and *HLA-DPB1* genes and discovered the most significant association with *HLA-DQB1*06:02*. Similarly, our lead variant rs9273375 was in high linkage disequilibrium (LD) with *HLA-DQB1*06:02* ($r^2$ = 0.888). As *HLA-DQB1*06:02* is also in high LD with *DRB1*15:01*, we estimated the pairwise LD also for *HLA-DRB1*15:01*. The analysis supported *HLA-DQB1*06:02* as the most significant HLA-allele to associate with Lyme disease.

**Supplementary Table 3.** *Linkage disequilibrium between lead SNP at the HLA-alleles.*

| Variant 1 | Variant 2 | $r^2$, D’ |
|---|---|---|
| rs9273375 | *HLA-DQB1*06:02* | 0.888, 0.990 |
| rs9273375 | *HLA-DRB1*15:01* | 0.877, 0.979 |
| *HLA-DRB1*15:01* | *HLA-DQB1*06:02* | 0.983, 0.997 |

*Variant 1 and Variant 2 represent single variants and HLA alleles for LD estimate. We represent LD as $r^2$ and D’ for each comparison using the genotype data and imputed HLA alleles from FinnGen.*

#### b) *SCGB1D2* locus

Our GWAS result showed a novel genome-wide significant finding related to Lyme disease in chromosome 11. We found that the lead intronic variant (rs4110197) in *SCGB1D2* was in high LD with a missense variant (rs2232950). This causes amino acid change from proline to leucine indicating a deleterious mutation by several algorithms [12] [13].

To examine *SCGB1D2* locus and its genomic variation's causality to Lyme disease in more detail, we finemapped this region utilizing the "Sum of Single Effects" -model, called *SuSiE* [11]. The 99% credible set included eight variants including the common missense variant rs2232950. Furthermore, *SuSiE* predicted posterior probability of 0.22 to the lead variant rs4110197.

### 4. PheWAS analysis

#### a) Comparison with publicly available traits

We performed a phenome-wide association analysis (PheWAS) to explore the association between the missense variant rs2232950 and 2,202 disease endpoints from FinnGen. FinnGen endpoints include primarily electronic health record derived phenotypes. To complement this analysis, we computed PheWAS also using the OpenTargets platform, which includes traits from publicly available GWASes and traits from other biobanks. This analysis did not reveal additional significant associations besides its association with Lyme disease in FinnGen and rs2232950. Therefore, this analysis did not provide additional insight into the function of *SCGB1D2* (Supplementary Figure 1).

### Supplementary Figure 1. Open targets PheWAS with rs2232950

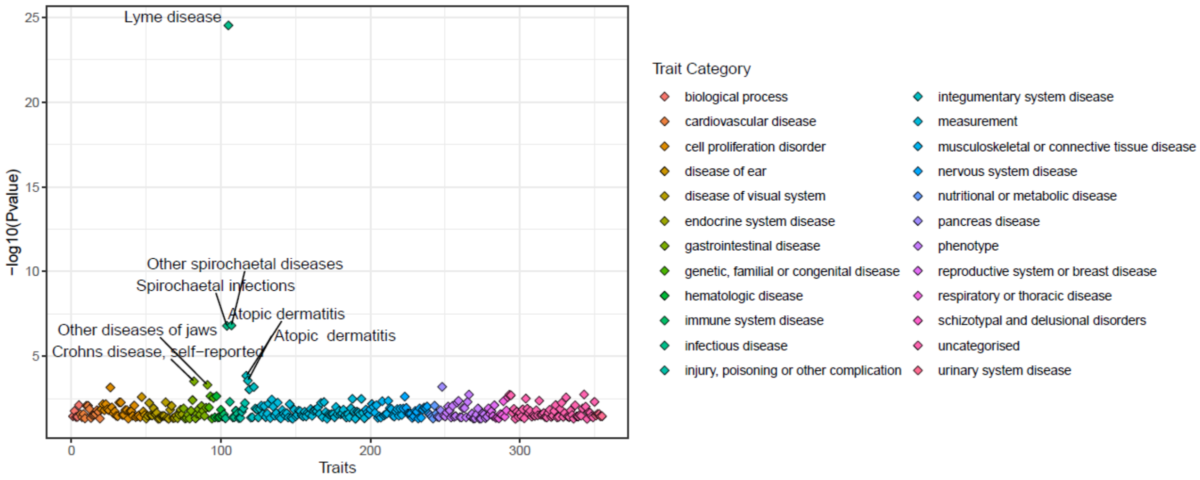


**Supplementary Figure 1.** *Phenome-wide association (PheWAS) from publicly available data from the OpenTargets platform are visualized by trait and their -log10(P-values) in the y-axis. Colors represent trait categories.*

### b) GWAS for syphilis

We were curious to investigate whether we could find associations between our findings in *SCGB1D2* and other fairly common spirochetal infections, such as syphilis. Therefore, we performed a GWAS for syphilis using FinnGen data with 719 individuals with syphilis ICD-codes captured from hospital inpatient, outpatient and primary care registries (Supplementary Figure 1). We observed a significant association with the *HLA-DRB5* locus but did not find a significant association between *SCGB1D2* locus and syphilis at single variant or locus level (rs2232950, P=0.6). This finding further supported that rs2232950 may be a specific predisposing component to Lyme disease.

Supplementary Figure 2. Analysis of *SCGB1D2* locus in syphilis

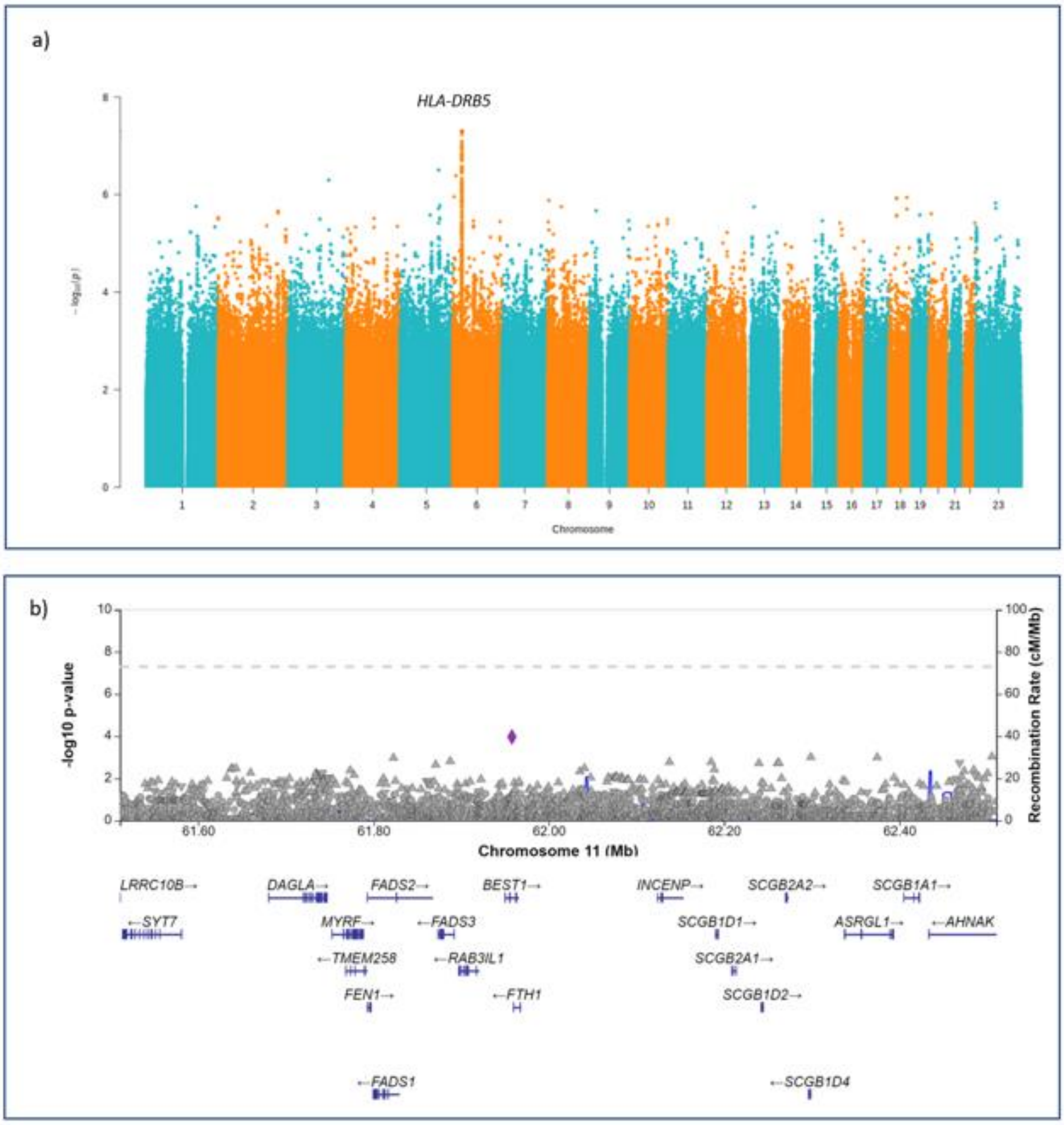


**Supplementary Figure 2.** ***a)*** *Manhattan plot for the genome-wide association study (GWAS) for syphilis including 719 individuals with syphilis diagnosis and 341,780 controls. HLA-DRB5 association seen also with syphilis is different from our HLA association in Lyme disease, and the P-value from the lead variant from Lyme disease HLA lead variant rs9273375 is P=0.45.* ***b)*** *Regional association of SCGB1D2 locus in syphilis. The variant with lowest P-value is marked here with a diamond. P-value for Lyme disease variant rs2232950 is not significant (P=0.6).*

### 5. Expression profiling

#### a) GTEx

We examined RNA expression across tissue types using The Genotype-Tissue Expression (GTEx) v8 using the fully processed and normalized gene expression matrices for each tissue. These data contain RNA expression samples from 948 donors across 54 tissues. These same values are used for eQTL calculations by GTEx. We extracted the values for *SCGB1D2* in these data and plotted the normalized values per tissue (Supplementary Figure 3). The two skin types (sun exposed and unexposed) show the highest expression levels of *SCBG1D2*.

Supplementary Figure 3. Tissue distribution of *SCGB1D2* expression

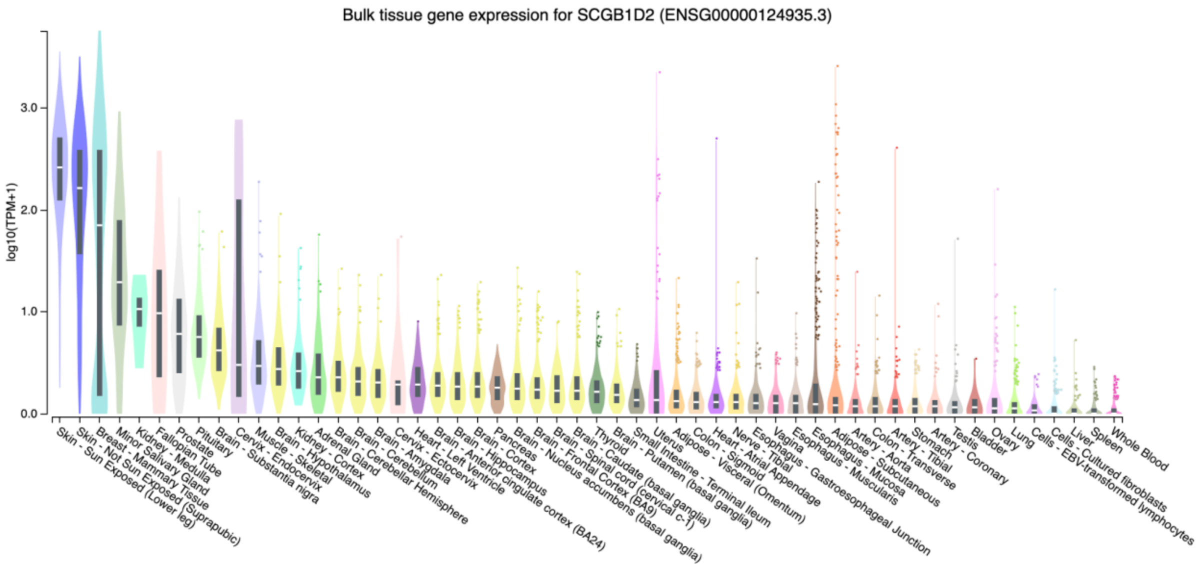


**Supplementary Figure 3.** *Expression pattern of SCGB1D2 across human tissues. We obtained RNA expression data from the GTEx project[16] and examined the expression profile of SCGB1D2 across tissues. A total of 701 individuals with expression values showed the highest expression in the skin (sun exposed) and skin (not sun exposed).*

#### b) Single cell analysis

In order to understand the relevant cell types for *SCGB1D2* expression from the skin we examined single cell sequencing data from skin. We observed that *SCGB1D2* was predominantly expressed by the sweat gland cells (Supplementary Table 4).

Supplementary figure 4. Expression of *SCGB1D2* by cell type in the skin

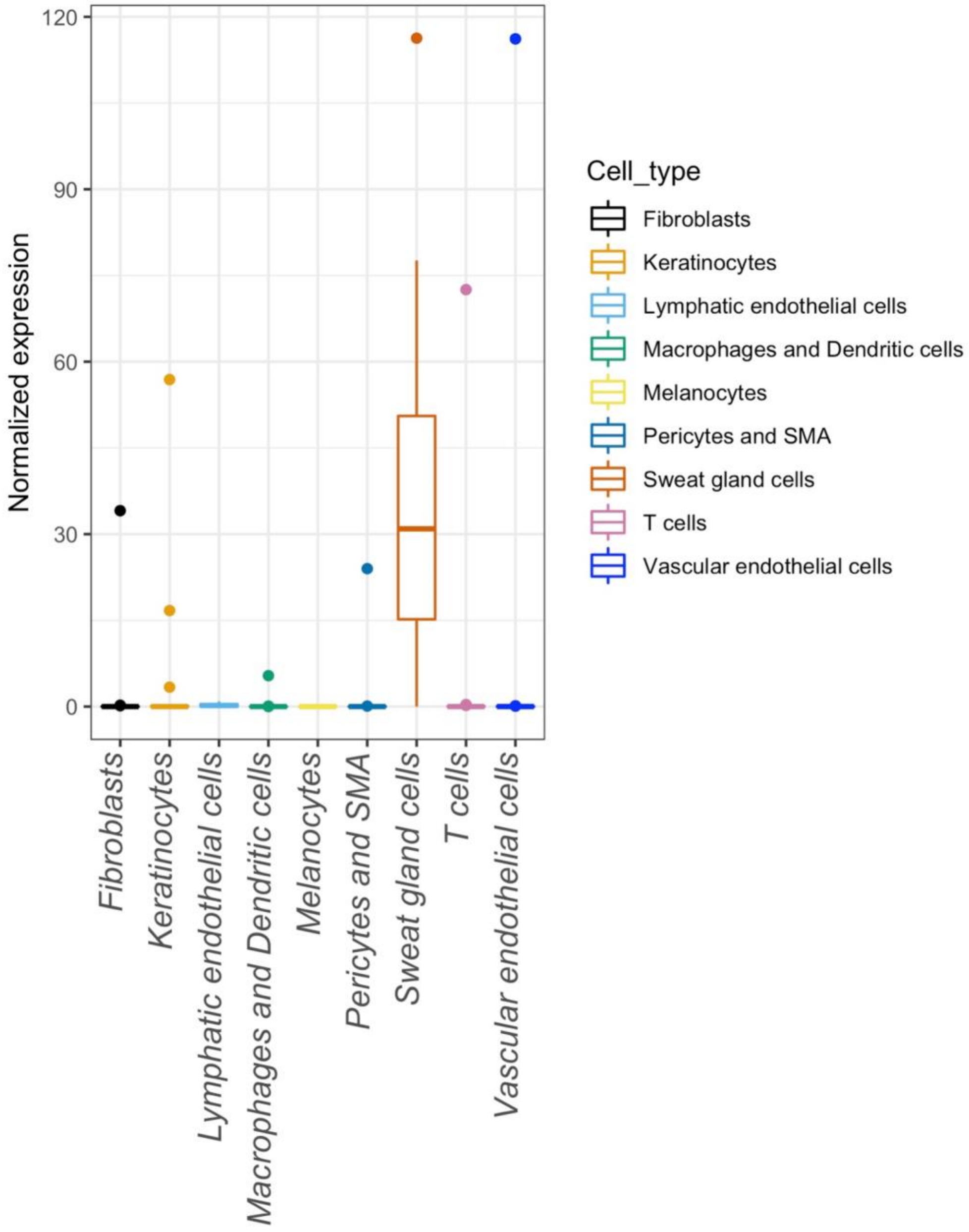


**Supplementary Figure 4.** *Single cell sequencing data shows SCGB1D2 expression is specific for sweat gland cells. Data from He et al.* [17]

### 6. Functional analyses of *Borrelia burgdorferi*

The results below support our main findings in live *Bb*. We estimate the effect of *SCGB1D2* P53L on *Bb* growth and examine the killing capability of *SCGB1D2* recombinant protein on live *Bb*.

#### Supplementary Figure 5. *Borrelia burgdorferi* growth inhibition by *SCGB1D2* P53L

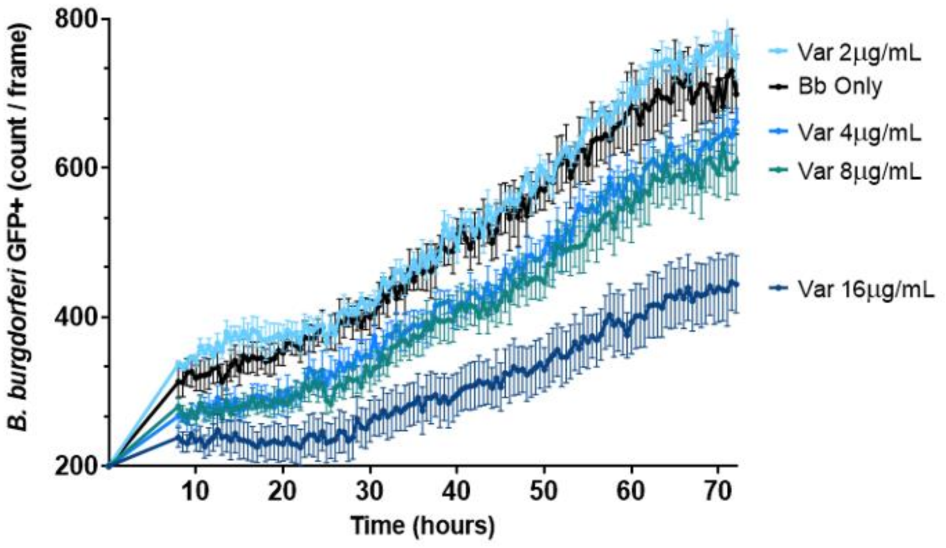


**Supplementary Figure 5.** *Timescale analysis of Borrelia burgdorferi growth inhibition by SCGB1D2* P53L *over 72h hours. Y-axis represents green fluorescent protein (GFP) count per frame and X-axis represents time. Concentrations tested are 2 to 16 µg/mL.*

Supplementary Figure 6. *Borrelia burgdorferi* killing by *SCGB1D2*

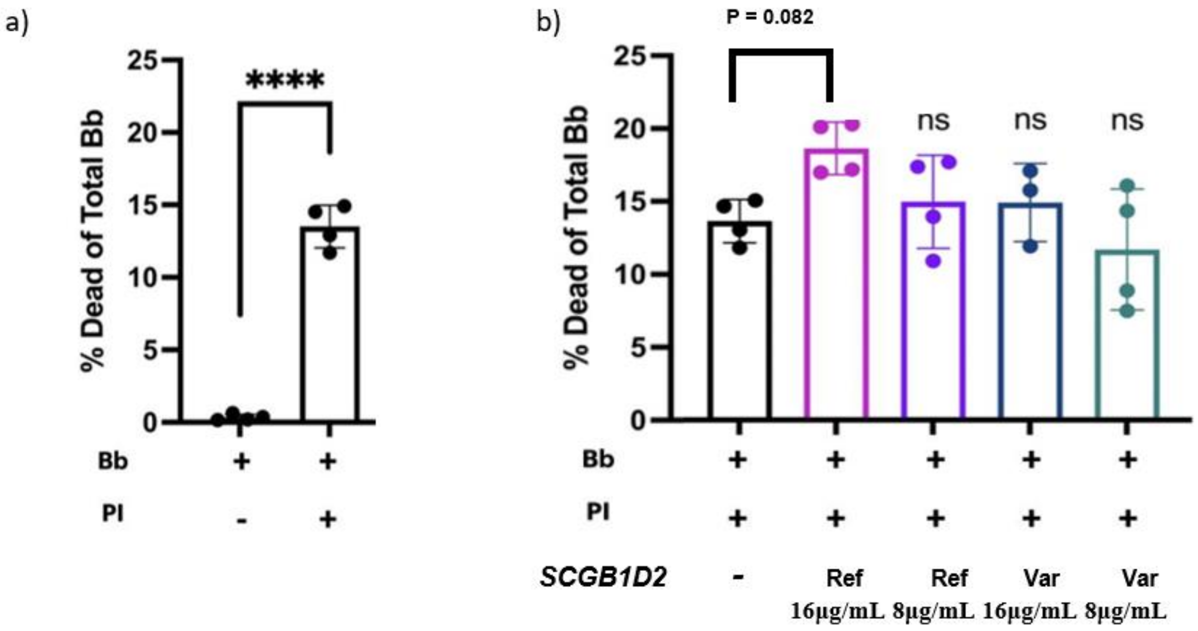


**Supplementary Figure 6.** *Borrelia burgdorferi (Bb) spirochetes expressing green fluorescent protein (GFP) were incubated with* ***a)*** *with or without propidium iodide (PI).* ***b)*** *either 8µg/mL or 16µg/mL of reference (Ref) or variant (Var) SCGB1D2 protein in the presence of PI to measure Bb death by SCGB1D2. After 24 hours of incubation, an aliquot of each culture was analyzed by flow cytometry. Overall one-way ANOVA was used comparing Bb with PI and Bb with PI and SCGB1D2 proteins (ANOVA F(4, 14 )=3.185, P=0.0467, Dunnett's multiple comparisons test P=0.082). ****P<0.0001; ns, not significant.*
